# Corticospinal excitability during timed interception depends on the speed of the moving target

**DOI:** 10.1101/2025.03.25.645232

**Authors:** Justin R. McCurdy, Daniel Zlatopolsky, Ria Doshi, Jing Xu, Deborah A. Barany

## Abstract

Successfully intercepting a moving object requires precisely timing the optimal moment to act by integrating information about the target’s visual motion properties. Neurophysiological evidence indicates that activity in the primary motor cortex (M1) during interception preparation is sensitive to both the target’s kinematic features and motor planning. However, how visual motion signals are integrated within M1 to guide interception timing is unclear. In the present study, we applied single-pulse transcranial magnetic stimulation (TMS) over M1 to examine how a target’s kinematics influence corticospinal excitability during interception preparation. Participants were instructed to abduct their right index finger to intercept a target moving horizontally at a constant speed toward a fixed interception zone. Target speed (Fast or Slow) and travel distance (Far or Close) were manipulated while controlling motion duration across conditions. Motor-evoked potentials (MEPs) were elicited at five latencies before target arrival at the interception zone. Consistent with previous behavioral findings, movement initiation occurred earlier for faster targets and was delayed when TMS was applied closer to the target’s arrival. Though MEPs were generally suppressed relative to baseline at earlier timepoints and facilitated closer to movement initiation, we observed that target speed—but not distance—influenced the time course of MEP modulation. When adjusting for movement initiation times, there was an overall reduced suppression and increased facilitation for faster-moving targets, possibly reflecting a heightened urgency to move. These results suggest M1 activity during interception preparation integrates internal estimates of target motion, which may serve to optimize interception timing and performance.

**New & Noteworthy:** When intercepting a moving object, like catching a ball, we need to continuously combine visual motion signals to predict the object’s future location and enable accurate movement. Here, we show that preparatory suppression and facilitation of corticospinal excitability depends on the speed, but not the distance, of the moving target. These findings reveal that differences in interception timing are closely linked to changes in motor system excitability.

## Introduction

Intercepting moving objects with the hand, like catching a ball or stopping a water bottle from rolling off the table, relies on integrating a continuous prediction of the object’s future location with estimates of the current hand position (Brenner and Smeets 2018; Crowe et al. 2023; De La Malla and López-Moliner 2015). Humans achieve high spatial and temporal precision during interceptive movements in part by estimating time-to-contact (TTC), defined as the time remaining until a moving object or intercepting effector reaches a specific location (Lee 1976; Senot et al. 2003; Merchant et al. 2009; Baurès et al. 2021). When intercepting objects moving at constant speed, humans use both kinematic (e.g., speed and distance) and timing cues to accurately estimate TTC (Chang and Jazayeri 2018). Under full vision conditions, movements are initiated earlier and movement duration is shorter for faster moving targets (Brouwer et al. 2000; Dubrowski et al. 2000), implying that accurate integration of kinematic information is crucial for optimizing interception timing (Brouwer et al. 2003; Senot et al. 2003; Nelson et al. 2019; Zhao et al. 2019).

The neural substrates responsible for estimating TTC to guide interceptive timing remain largely unexplored (Fooken et al. 2023). In addition to engaging visuomotor fronto-parietal networks (Battaglia-Mayer and Caminiti 2018; Gallivan and Culham 2015), estimating TTC while preparing to intercept a moving target is associated with neural activity in the middle temporal area (MT/V5+), sub-cortical regions, and the cerebellum (Senot et al. 2008; Baurès et al. 2021; De Azevedo Neto and Bartels 2021; Polti et al. 2022; Li et al. 2015; Bosco et al. 2008; O’Reilly et al. 2008). Most of these signals are expected to converge within primary motor cortex (M1) to specify and release descending motor commands (Hanes and Schall 1996). Evidence from single-neuron recordings in non-human primates and human neuroimaging indicate that the modulation of M1 activity reflects both time-varying aspects of motor planning and processing of visual information to estimate TTC for interception (Daneshi et al. 2020; Field and Wann 2005; Merchant et al. 2009; Port et al. 2001; Senot et al. 2008).

In humans, single-pulse transcranial magnetic stimulation (TMS) has been extensively used to study corticospinal excitability (CSE) modulation during action preparation (reviewed in Bestmann and Duque 2016; Derosiere et al. 2020). In delayed reaction time (RT) and predictive timing tasks, in which a cue specifying an upcoming movement is followed by an external or internal imperative signal to move, a consistent pattern of CSE modulation prior to movement onset is observed. Typically, CSE is suppressed relative to baseline, followed by rapid facilitation in the agonist muscle, coupled with increasing suppression in the antagonist muscle (Ibáñez et al. 2020; Badry et al. 2009; Duque et al. 2017; Duque and Ivry 2009; Greenhouse et al. 2015; Hasbroucq et al. 1997; Labruna et al. 2014, 2019; Lebon et al. 2016; Touge et al. 1998; Vassiliadis et al. 2020). In addition, the TMS pulse itself affects timing of the motor response: in simple RT tasks, TMS applied prior to onset can shorten RT, whereas in predictive timing tasks TMS delays movement onsets (Ibáñez et al. 2020; Leocani et al. 2000; Nickerson 1973; Pascual-Leone et al. 1992). Though the functional significance of these TMS-elicited changes in CSE and movement is unresolved, it is thought to reflect a transitional state that primes the motor system for voluntary movement, involving preparatory changes in CSE that ready muscles to initiate the upcoming action (Haith and Bestmann 2020; Ibáñez et al. 2020; Quoilin et al. 2019).

While previous studies have explored CSE during anticipatory actions in response to moving stimuli (Coxon et al. 2006; Ibáñez et al. 2020; Marinovic et al. 2009, 2010), it is unclear how systematically varying kinematic motion properties may affect CSE during motor preparation for interception. In this study, we investigated how two TTC variables – target speed and distance – affect CSE when preparing to intercept moving targets with the hand. We measured TMS-elicited motor-evoked potentials (MEPs) in the task-relevant right first dorsal interosseous (FDI) and task-irrelevant abductor digiti minimi (ADM) muscles as participants prepared to abduct their index finger to coincide with a moving target’s arrival at a fixed interception location. On each trial, we varied the target’s speed (Fast or Slow) and starting distance (Close or Far), along with the timing of the TMS pulse relative to the ideal interception time. We hypothesized that faster targets and closer starting distances would lead to increased modulation of excitability due to heightened response urgency. Additionally, we hypothesized that if M1 is involved in integrating target motion kinematics to estimate TTC, then the time course of CSE suppression and facilitation would align with interception onset.

## Materials and Methods

### Participants

We recruited 23 healthy, right-handed participants (12 M, 21.8 ± 3.6 years) with no known history of neurological disorders, no contraindications to TMS, and normal or corrected-to-normal vision to complete the experiment. All participants were naïve to the purpose of the study, provided written informed consent, and were compensated for their time. One participant was excluded for poor behavioral performance (see *Data Analysis*), resulting in a final sample of 22 participants (11 M, mean age 21.9 ± 3.7 years). The experimental procedures were approved by the Institutional Review Board at the University of Georgia.

### Experimental Setup

Participants were seated in front of the task monitor (1920 × 1080 resolution, 60 Hz refresh rate) at a desk with their right hand resting on a digitizing trackpad (WACOM Intuous Pro Large). The experiment and stimulus presentation were controlled using PsychoPy 3 Version 2022.2.4 (Peirce et al. 2019). The trackpad was positioned so that participants could rest both arms comfortably on the desk, while placing their chin in a custom stand approximately 58 cm from the center of the screen. Participants wore earplugs for the duration of the study and a sleeve over their index finger to enhance trackpad sensitivity. Surface electromyography (EMG) electrodes were attached to the FDI and ADM muscles of the participant’s right hand. EMG activity was recorded using a Delsys Bagnoli™ desktop system (Delsys Inc., Boston, MA, USA). EMG signals were amplified 1000x, bandpass filtered (20-450 Hz), and sampled at 5000 Hz through a data acquisition board (Micro 1401-4, Cambridge Electronic Design, Cambridge, UK) and CED Signal software.

### Transcranial Magnetic Stimulation

TMS was applied through a 70-mm, figure-of-eight coil driven by a DuoMAG MP magnetic stimulator (Deymed Diagnostics sro, Hronov, Czech Republic). MEPs were elicited from the right FDI by applying TMS over the hand area of the left M1. The coil was placed tangentially on the scalp with the handle oriented towards the occiput and laterally from the midline at a 45_°_ angle. Before the experimental session, the FDI motor hotspot (i.e., the optimal location for eliciting MEPs in the target muscle) was identified on each participant by eliciting TMS every 4 s and systemically repositioning the coil. At the beginning of each experimental session, a model of the brain was scaled to the participant’s cranial dimensions using a neuronavigational tracking system (BrainSight, Rogue Research, Montreal, Canada) to optimize and replicate coil position throughout the session. Resting motor threshold (rMT) was defined as the minimal TMS intensity required to evoke peak-to-peak MEP amplitudes of ∼50 uV in the target muscle for 5 of 10 consecutive trials (Rossini et al. 1994). The average rMT for this experiment was 43.3% (±4.5) of the maximum stimulator output. Stimulator intensity during the task blocks was set to 115% of participant’s rMT. TMS timings were controlled by sending TTL signals from the task computer to the stimulator through CED Signal.

### Interception Task

Participants performed an interception task in which the goal was to move an on-screen cursor representing their right index finger position into a pre-defined interception zone (IZ) timed to match the arrival of a moving target (Figure 1A). At the start of each trial, participants were instructed to move an on-screen cursor (black cross, 1 x 1 cm) into a fixed start position on the trackpad and remain there until the target appeared. Once the cursor was in the start position for 500 ms, a red rectangular target (0.65 x 4 cm) would appear on the screen at a fixed distance to the left of the IZ (0.65 x 4 cm). After a variable delay (500 - 2000 ms, uniform distribution), the target would begin approaching the interception zone at a constant horizontal speed. Participants were instructed to make a quick swiping movement by abducting their right index finger to intercept the target as it arrived at the IZ. During practice, participants were reminded to only move their index finger while keeping all other fingers at rest.

**Figure 1.**
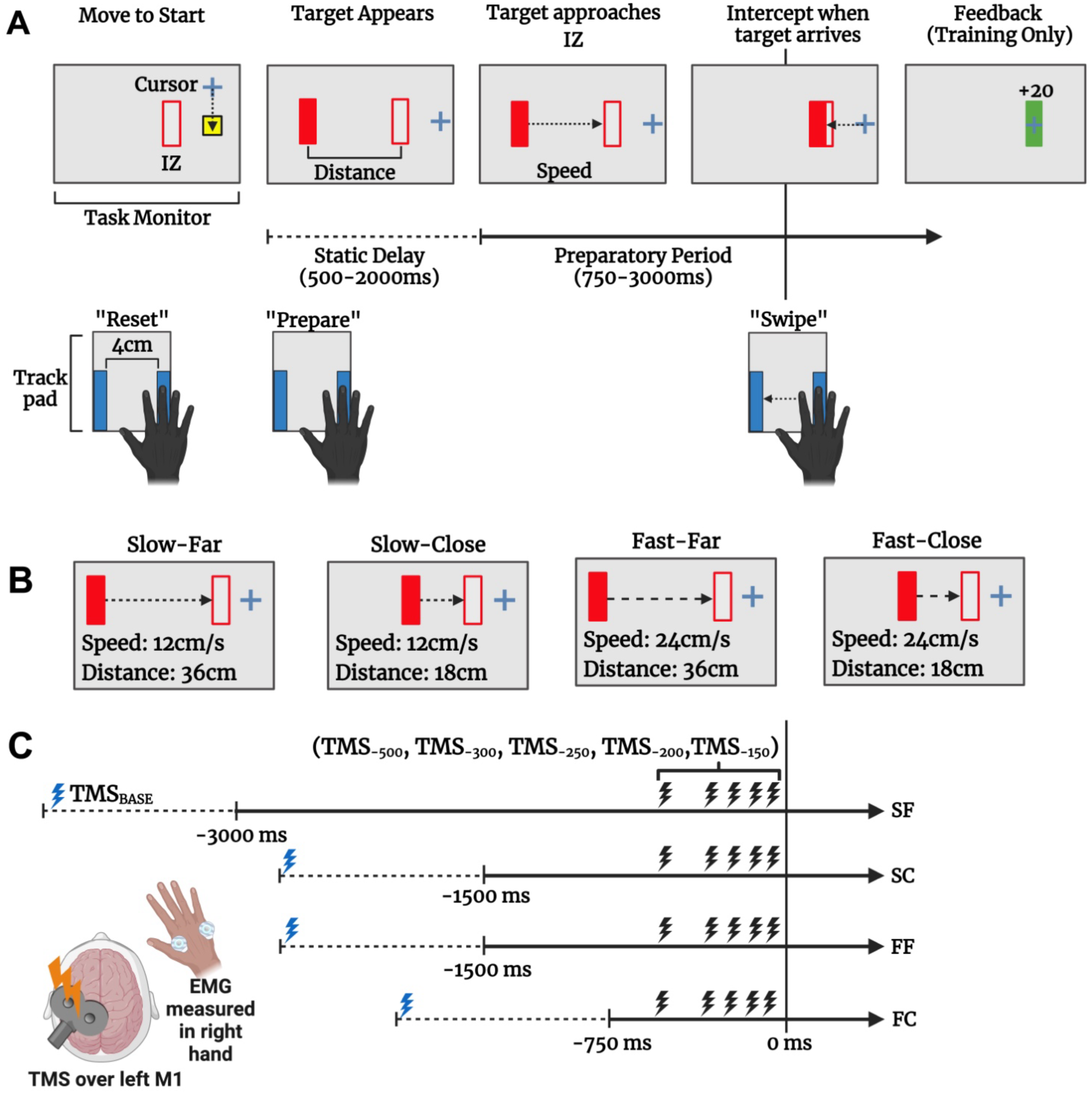
Interception task design. (**A**) Participants prepared to swipe their right index finger on a trackpad to intercept a moving target to coincide with the target time of arrival at a pre-defined interception zone (IZ) on the screen. During training, they were provided feedback about the movement timing. (**B**) During each block, the four trial conditions (Slow-Far, Slow-Close, Fast-Far, Fast-Close) were presented in randomized order, so participants needed to judge target motion to time their movements. Note that the four combinations of target speed and distance resulted in 3 unique preparation period durations (3000 ms for Slow-Far, 1500 ms for Slow-Close and Fast-Far, and 750 ms for Fast-Close). (**C**) TMS was applied over the left M1 representation of the first dorsal interosseous (FDI) muscle at target presentation (baseline) or at 1 of 5 time points (500, -300, -250, -200, or -150 ms) relative to target-IZ overlap during the preparation period. EMG was measured in both the task-relevant FDI muscle and the task-irrelevant abductor digit minimi (ADM) muscle.

On each trial, the target started at either 36 cm (Far) or 18 cm (Close) from the IZ and approached the IZ at either 12 cm/s (Slow) or 24 cm/s (Fast). Figure 1B depicts the 4 unique trial conditions based on the target’s visual motion properties: Fast-Close, Fast-Far, Slow-Close, and Slow-Far. Target motion durations were 750 ms for Fast-Close, 3000 ms for Slow-Far, and matched at 1500 ms for the Fast-Far and Slow-Close conditions to test for the effect of target kinematics independent of preparation time (Lebon et al. 2016). After 20 familiarization trials, participants performed two training blocks of 52 trials in which no TMS was applied. During training, feedback about the distance of the target from IZ at time of interception was provided to improve response timing.

After both training blocks were complete, participants performed 6 task blocks of 48 trials each. During task blocks, both the trial condition and time of TMS delivery were pseudo-randomized based on the block order assigned to the participant. During 44 of the 48 trials in each task block, TMS was applied at either the start of the trial to measure within-task baseline (TMS_BASE_; n=4 per block) or at one of five time points relative to the target’s arrival (i.e., 500, 300, 250, 200, or 150 milliseconds before target-IZ overlap; n=8 each per block). This resulted in a total of 24 in-task baseline TMS trials and 12 TMS trials per unique trial condition by time point combination. In the remaining 4 of the 48 trials in each task block (24 total trials), no TMS was applied to minimize expectancy effects (Tran et al. 2021).

### Data Analysis

One participant was excluded from data analysis for low interception accuracy (<50% of targets intercepted within IZ). Dependent variables were extracted from the EMG recordings and the finger position data on the trackpad. All EMG recordings were analyzed offline using the MATLAB VETA toolbox (Jackson and Greenhouse 2019). Interception performance was quantified by movement onset, movement time, and spatial accuracy. Movement onset was identified as the first timepoint the EMG activity in the FDI muscle exceeded 3 standard deviations of the mean of the rectified signal for the entire trial epoch and was >0.5 mV, relative to the target’s arrival time. Movement time was defined as the duration from EMG-derived movement onset to the time at which the IZ was reached. Spatial error was defined as the absolute value of the relative distance between the target and IZ at time the finger position entered the IZ.

CSE was indexed by the TMS-elicited MEP peak-to-peak amplitude. The root mean square (RMS) value of the background EMG activity was calculated for the 300 ms before the TMS artifact. Trials were rejected from analysis when the background RMS was □2 standard deviations from the participant’s average RMS (1.7% of all trials). Trials in which EMG activity was observed within 300 ms of the MEP and trials in which the EMG burst onset preceded the TMS pulse were excluded from analysis (0.8% of all trials). These steps were performed to prevent the contamination of the MEP measurements from fluctuations in background EMG activity (Darling et al. 2006). Lastly, trials in which the EMG burst onset were recorded more than 300 ms before or after the target’s TTC were removed to account for behavioral outliers (0.7% of all trials). Altogether, a total of 165 trials (3.1% of total trials) across all participants were excluded from this dataset. The peak-to-peak amplitude of the EMG signal was calculated during the 20-50 ms window after TMS (i.e., the normal range at which an MEP is expected to occur). Each participant’s MEP values were normalized to the participant’s average baseline MEP amplitude.

### Statistical Analysis

Interception performance in blocks without TMS was assessed using either a 2-way repeated measures ANOVA, with target speed (Fast or Slow) and starting distance (Close or Far) as within-subjects’ factors, or a one-way repeated measures ANOVA with motion duration (0.75 s, 1.5 s, 3.0 s) as the within-subjects factor.

Interception performance in TMS blocks and MEP peak-to-peak amplitude for the FDI and ADM muscles were assessed using a 3-way repeated measures ANOVA (2 speeds x 2 distances x 5 TMS timepoints). Additionally, for each TMS timepoint, planned one-sample t-tests compared changes in interception performance to no-TMS blocks and assessed whether MEP amplitude differed from baseline. Planned paired t-tests compared MEP amplitudes between FDI and ADM muscles at each TMS timepoint. For all ANOVA comparisons, the alpha level was set to 0.05. Effect sizes are reported using generalized η², and relevant post-hoc comparisons were conducted using the Holm correction (Holm 1979).

To further examine the interaction between target speed and time of TMS delivery for FDI, we performed a growth curve analysis (Wiley 2014) averaged over the levels of target speed. The overall time course of MEPs was modeled with a second-order (quadratic) orthogonal polynomial, including fixed effects of target speed (Fast vs. Slow; within-participants) on all time terms. The model also included participant random effects on all time terms and participant-by-condition random effects on all time terms. Parameter estimate degrees of freedom and corresponding p-values were estimated using Satterthwaite’s method (Satterthwaite 1946).

## Results

### Fast target speeds resulted in earlier movement onsets and decreased spatial accuracy

To assess performance differences driven by the target’s motion properties independent of stimulation, behavioral data were first evaluated from the last block of trials without TMS (Figure 2). Across all trial conditions, movements were initiated on average 91.7 ms (±24.4) prior to the target’s arrival (Figure 2A). A 2-way repeated-measures ANOVA (2 speed x 2 distance) showed significant effects of both target speed (*F*_1,21_=46.76, *η*^2^=.128, *p*<0.001) and start distance (*F*_1, 21_ = 5.44, *η*^2^=.026, *p*=0.03), indicating that participants-initiated movement earlier for faster-moving targets and for targets starting farther away. There was also a significant two-way interaction (*F*_1, 21_ = 7.00, *η*^2^=.021, *p*=0.015), indicating that movement onsets were earlier for faster moving targets only at farther starting distances. Post-hoc tests showed significant differences between Fast-Close and Fast-Far (*t* (21) =3.31, *p*=0.01), Slow-Close and Fast-Far (*t* (21) =4.61, *p*<0.001), and Slow-Far compared to Fast-Far (*t* (21) =-4.48, *p*=0.001), suggesting that faster targets induced earlier movement onsets even with the same motion duration.

**Figure 2.**
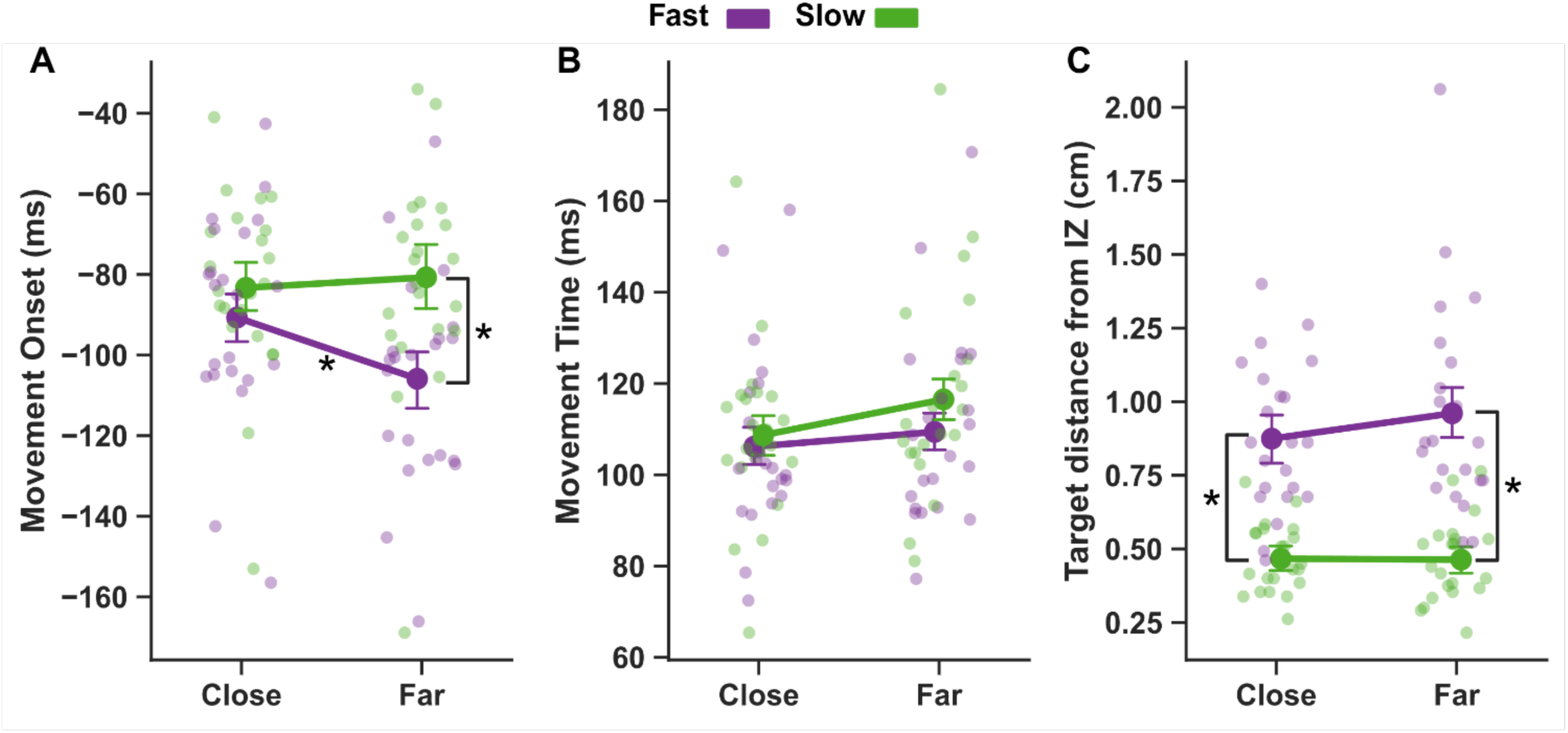
Target speed and preparation duration affected interception characteristics. (A) Movement onset (ms) was earlier for Fast targets at Far starting distances. (B) Movement time (ms) was significantly faster for Fast-Close (750 ms duration) than Slow-Far (3000 ms duration). (C) Spatial error (cm) was higher in Fast target conditions. Error bars correspond to bootstrapped 95% confidence interval. \**p* <.05 between conditions.

Movement times (See Figure 2B) were significantly faster when the target started from the close position (*F*_1, 21_= 9.82, *η*^2^=.022, *p*=0.005) and when approaching at higher speed (*F*_1, 21_= 7.30, *η*^2^=.019, *p*=0.013). There was no significant interaction between target speed and distance (*F*_1, 21_= 1.31, *η*^2^=.006, *p*=0.265). Post-hoc tests showed movement times were significantly different between the Fast-Close and Slow-Far condition (*t*(21) = 3.67, *p* < 0.013), but not between any other condition pairs (all *p’s* > 0.05). In addition, grouping the movement times based on motion duration (Fast-Close; 750 ms, Fast-Far and Slow-Close; 1500 ms, Slow-Far; 3000 ms) revealed a significant main effect (*F*_1.64, 34.40_=7.64, *η*^2^=.055, *p*=0.003). Post-hoc tests revealed a significant difference between the 750 and 3000 ms durations (*t* (21) = -3.66, *p*=0.004), but not between the 750 and 1500 ms (*t* (21) = -1.23, *p*=0.234), or the 1500 ms and 3000 ms durations (*t* (21) = -2.41, *p*=0.051). This suggests that movement times were likely influenced mainly by the duration of the preparatory period.

Participants intercepted 76% (± 8.7) of targets within the IZ, indicating overall accurate task performance (Figure 2C). To more closely examine interception accuracy, we calculated spatial error on each trial as the average distance of the target to IZ at the time of interception. Spatial errors were larger for fast compared to slow targets (*F*_1,21_= 104.8, *η*^2^=.494, *p*<0.001), but did not differ across levels of start distance (*F*_1,21_=2.07, *η*^2^=.013, *p*=0.165). The interaction effect was not significant (*F*_1,21_=1.76, *η*^2^=.012, *p*=0.199). Together, these results indicate that interception accuracy decreased at faster target speeds.

### TMS timing differentially altered interception performance by target speed and distance

To examine the influence of applying TMS on interception performance, the average of each subject’s movement onset, movement time, and interception accuracy were calculated for each combination of target condition (Fast-Close, Fast-Far, Slow-Close, Slow-Far) and TMS delivery time (500, 300, 250, 200, 150 ms prior to target’s arrival) and then normalized to each subject’s average during the second block of trials without TMS (Figure 3).

**Figure 3.**
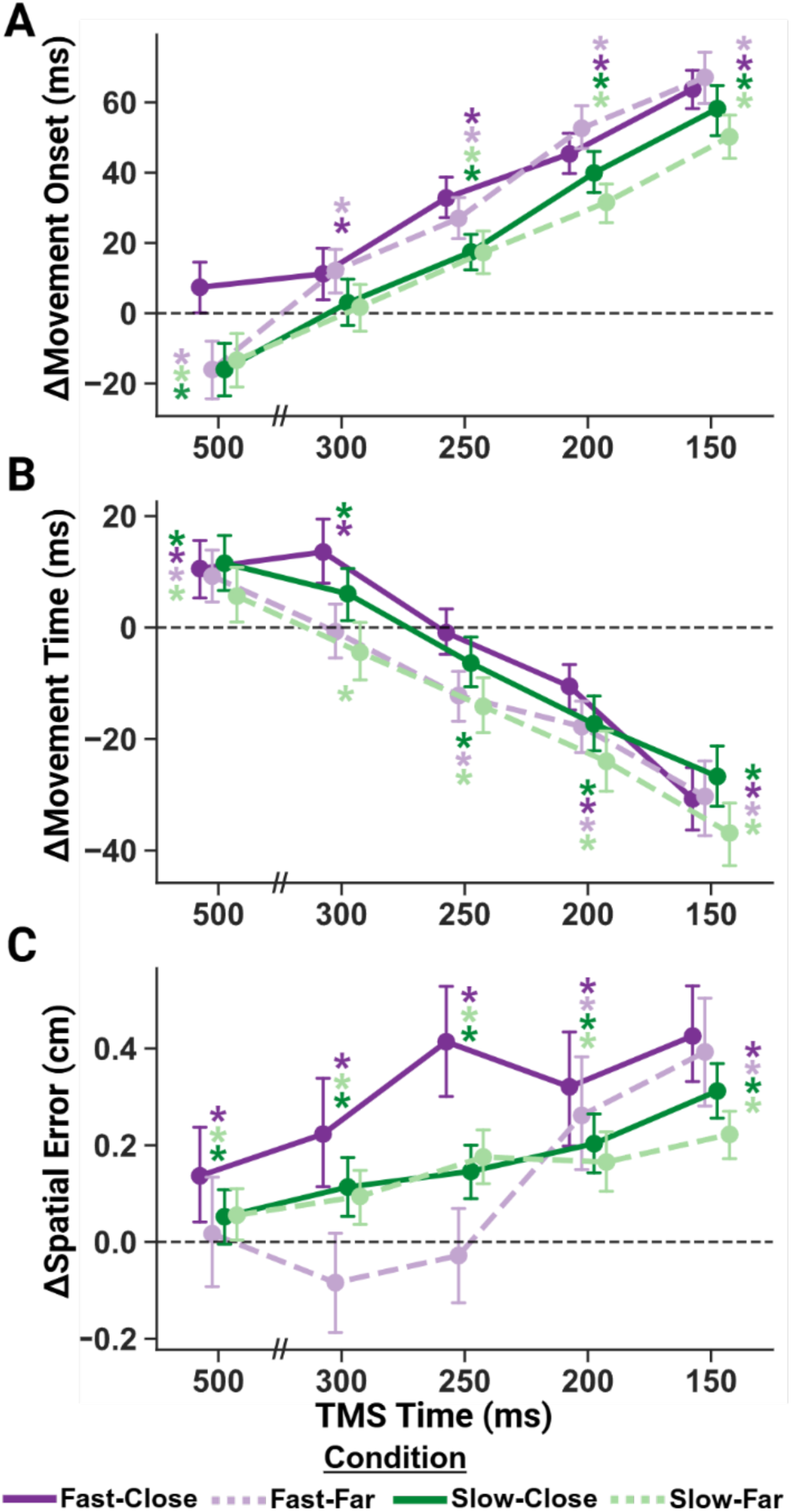
TMS applied during movement preparation disrupts interception timing. All plots show average change relative to the participant’s average on trials without TMS. Purple (green) lines represent Fast (Slow) target trials and solid (dotted) lines represent Close (Far) start distance trials. (**A**) Movement onset was delayed more at TMS time points closer to the target arrival and for Fast target conditions. Circles show the average for each condition at each time of TMS delivery. (**B**) Movement time sped up at TMS time points closer to target arrival and for Far target conditions. (**C**) Spatial error tended to increase during TMS trials, especially at later time points. All error bars correspond to bootstrapped 95% confidence interval. \**p* <.05 different from zero.

For change in movement onset (Figure 3A), there was a significant main effect of time of TMS delivery (*F*_1.78, 37.47_=79.20, *η*^2^=0.440, *p*<0.001). In general, TMS delayed movement onset more when applied closer to the target’s time of arrival (all *p’*s < 0.05), suggesting that TMS during late stages of movement preparation interferes with response execution. There was a significant main effect of target speed (*F*_1,21_=15.06, *η*^2^=0.045, *p*<0.001), but not of start distance (*F*_1,21_=2.66, *η*^2^=0.006, *p*=0.188), indicating that the change in movement onset was greater for faster moving targets. There was a significant three-way interaction between target speed, distance, and time of TMS delivery (*F*_2.93, 61.43_=7.86, *η*^2^=0.013, *p*<0.001). Post-hoc tests revealed that at the earliest timepoint (500 ms before target arrival), TMS sped up movement onsets for the Fast-Far (*t* =-3.19, p=0.002), Slow-Close (*t* =-5.12, *p*<0.001), and Slow-Far (*t* =-3.44, *p*<0.001) conditions, but not for the Fast-Close (*t* =1.44, *p*=0.151) condition. In contrast, at 300 ms, stimulation delayed movement onset for the Fast-Close (*t* =3.28, *p*=0.001) and Fast-Far (*t* (258) =6.46, *p*<0.001) conditions, but not for the Slow-Close (*t* =0.937, *p*=0.35) or Slow-Far (*t* =1.66, *p*=0.099). Movement onset was significantly delayed for each condition at TMS_250_, TMS_200_, and TMS_150_ (all *p’s*<0.001). Together, these results suggest that TMS delays movement initiation more when applied closer to movement initiation and for faster-moving targets.

Analysis of movement times suggests that participants sped up or slowed down their movements to partially compensate for differences in movement onset (Figure 3B). There was a significant main effect of TMS stimulation time (F_1.56, 32.83_=36.79, *η*^2^=0.311, p<0.001) - movement times were shorter as TMS was delivered closer to target arrival time relative to blocks with no TMS. There was a significant main effect of target distance (*F*_1,21_=9.46, *η*^2^=0.034, *p*=0.006), but not target speed (F_1,21_=1.96, *η*^2^=0.006, p=0.177), indicating that TMS shortened movement times more for Far target distances. In addition, a significant two-way interaction between target distance and TMS time was observed (F_2.92_, *η*^2^=0.008_, 61.29_=3.78, *p*=0.02). When TMS was delivered 500 ms before target arrival, movement times were significantly increased for all conditions (all *p’s* < 0.05). At 300 ms prior to target arrival, movement times were significantly increased for the Fast-Close (*t* =4.35, *p*<0.001) and Slow-Close (*t* =3.96, *p*<0.001), decreased for Slow-Far (*t* = -3.28, *p*=0.001), and not significantly different from zero for Fast-Far (*t* =-0.278, *p*=0.781) trial conditions. At 250 ms before target arrival, TMS shortened movement times for both Far conditions (Fast-Far (*t* =-4.80, *p*<0.001); Slow-Far (*t* =-5.81, *p*<0.001)), but not the Close conditions (Slow-Close (*t* = -1.81, *p*=0.072); Fast-Close (*t* = -4.8, *p*=0.991). At both the latest TMS delivery timepoints (-200 and -150 ms), movement times were significantly shortened for all conditions (all *p’s* < 0.001). Altogether, TMS during earlier stages of movement preparation resulted in longer movement times, while TMS during late stages of movement preparation resulted in shorter movement times, especially when the target started from farther away.

TMS increased spatial error, especially when delivered closer to the target’s arrival (Fig. 3C). There was a significant main effect of TMS time (*F*_2.82, 56.48_=9.86, *η*^2^=.096, *p*<0.001) such that change in spatial error was greater at later TMS timepoints. There was a significant main effect of target distance (*F*_1,20_= 14.84, *η*^2^=.054, *p*<0.001), but not target speed (*F*_1,20_=2.29, *η*^2^=0.013, *p*=0.146), indicating increased spatial error for Close target distances. Furthermore, we observed a significant two-way interaction between target speed and distance (*F*_1,20_=14.59, *η*^2^=0.046, *p*<0.001) as well as a three-way interaction between target speed, distance, and TMS time (*F*_2.99, 59.88_=4.46, *η*^2^=0.019, p=0.007). When TMS was elicited at 200 or 150 ms prior to target arrival, stimulation increased spatial error (*p*<0.001, for all conditions). At earlier timepoints, TMS increased spatial error for all but the Fast-Far condition. Collectively, these results suggest that TMS interferes with the late stages of movement preparation to reduce interception accuracy, and that the ability to compensate for the effects of TMS is influenced by both target speed and distance.

### Corticospinal excitability depended on target speed, but not distance

To evaluate CSE modulation, we calculated changes in MEP amplitudes normalized to within-task baseline at each time of TMS delivery for the task-relevant FDI and task-irrelevant ADM muscles. Figure 4 shows the average normalized amplitude of MEPs evoked in FDI and ADM at each TMS time across all conditions. There were significant main effects of TMS time (F_1.87,39.29_=8.43, η^2^=0.055, *p*=0.001) and muscle (F_1,21_=12.1, η^2^=0.09, *p*=0.002), showing that overall excitability was significantly higher for FDI relative to ADM. A significant interaction (F_1.83,38.41_=19.1, η^2^=0.114, *p*<0.001) revealed that CSE was modulated differently throughout the preparatory period depending on the muscle’s relevance to the upcoming movement. Follow-up tests showed that for FDI, CSE was suppressed relative to baseline early in the preparatory period, followed by facilitation just prior to movement execution (*F*_1.79, 37.57_ = 14.97, η^2^=0.189, p<0.001). Mean FDI excitability was significantly decreased relative to baseline when TMS was elicited at 500 ms (*t* = -3.11, *p*=0.002), 300ms (*t* = -7.97, *p*<0.001), and 250 ms (*t*=-5.92, *p*<0.001), and increased to above the baseline at 150 ms prior to movement onset (*t*= 7.77, *p*<0.001).

**Figure 4.**
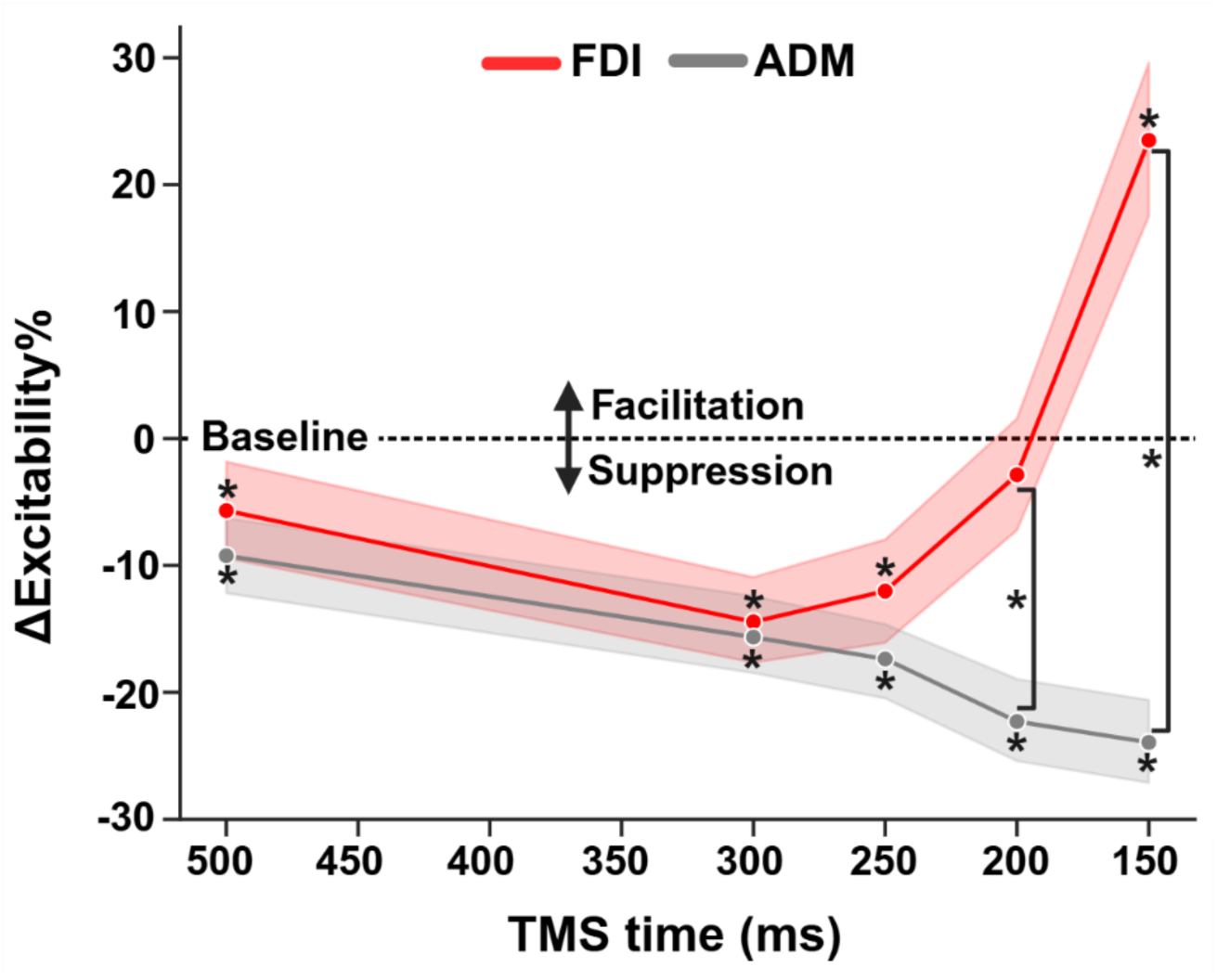
Early task-relevant suppression and late facilitation during interception preparation. Timeline of corticospinal excitability for task-relevant (FDI) and task-irrelevant (ADM) effectors across all target speed and distance conditions, showing early preparatory suppression and late facilitation. Error bars correspond to bootstrapped 95% confidence interval. \**p* <.05 different from zero or between FDI and ADM muscles.

The task-irrelevant muscle (ADM) showed suppression that increased as TMS was elicited closer to the target’s time of arrival (*F*_1.79, 37.57_=8.38, η^2^=0.07, *p*<0.001). Post-hoc tests confirmed that ADM excitability was significantly suppressed relative to baseline at all TMS timepoints (all *p’s* < 0.001). Pairwise t-tests between the muscle conditions at each level of

Excitability was significantly different between the FDI and ADM muscles when TMS was delivered 150 ms (*p* < 0.001) and 200 ms (*p <* 0.001) before target arrival, but not at any earlier TMS timing. Together, these results confirm that during movement preparation, task-irrelevant muscles undergo a sustained and increasing suppression (Sohn and Hallett 2004; Beck and Hallett 2011; Ibáñez et al. 2020).

Figure 5 shows the average change in CSE from within-task baseline at each TMS time point for each of the four target motion conditions. For the FDI muscle, we conducted a 3-way repeated measures ANOVA, including the target’s speed (Fast vs. Slow), start distance (Close vs. Far), and time of TMS relative to the target’s time of arrival (-500, -300, -250, -200, -150 ms) as factors. There was a significant interaction between TMS timepoint and target speed (*F*_2.38,50.23_=3.24, η^2^=0.010, *p*=0.039), but not distance (*F*_3.23, 67.83_=0.72, η^2^=0.002, *p*=0.556), suggesting that CSE was modulated throughout interception preparation differently depending on the speed of the target. In addition, main effects of target speed (*F*_1,21_= 4.93, η^2^=0.005, *p*=0.038) and TMS time (*F*_1.79, 37.63_=14.90, η^2^=0.142, *p*<0.001) were observed, indicating increased excitability for faster-moving targets.

**Figure 5.**
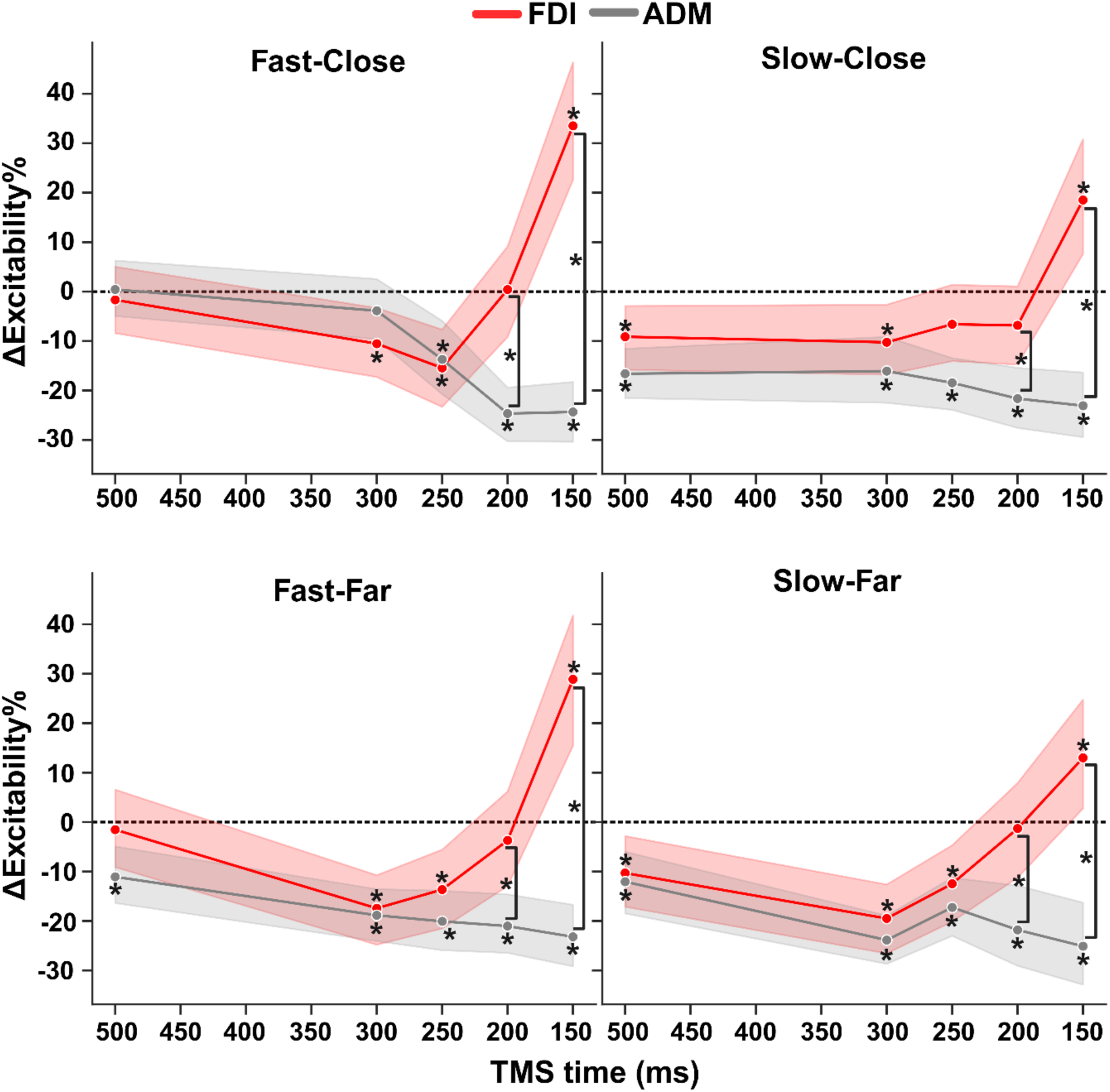
Corticospinal excitability (CSE) during interception preparation shows higher late facilitation for faster-moving targets and earlier suppression for slower-moving targets. Each subplot shows the timecourse of change in CSE relative to baseline for relevant (FDI) and nonrelevant (ADM) muscles for each target motion condition. Error bars correspond to bootstrapped 95% confidence interval. \**p* <0.05 from zero or between muscles.

Follow-up t-tests showed that when TMS was delivered at 150 ms before the target’s arrival at IZ, there was a significant increase in the MEP amplitude relative to baseline for all 4 motion conditions (Fast-Close; (*t* =5.41, *p*<0.001), Fast-Far; *t* = 4.38, *p*<0.001), Slow-Close; (*t* =3.31, *p*=0.001), Slow-Far; (*t* =2.28, *p*=0.02)). In contrast, at 200 ms before target TTC, MEP amplitudes for all 4 trial conditions were not significantly different from zero (all *p’*s > .05). MEPs elicited at 250 ms before target arrival were significantly reduced for Fast-Close (*t* =-3.88, *p*<0.001), Fast-Far (*t* =-3.41, *p*<0.001), and Slow-Far (*t* = -3.07, *p*=0.002) conditions, but not for Slow-Close (t = -1.57, *p*=0.117). MEPs elicited at 300 ms were significantly reduced for all trial conditions (Fast-Close; (*t* =-2.85 *p*=0.004), Fast-Far; *t* = -4.83, *p*<0.001), Slow-Close; (*t* = -2.74, *p*=0.007), Slow-Far; *t* =-5.67, *p*<0.001)). Finally, at our earliest TMS time point (500ms from target-IZ overlap), MEP amplitude was significantly reduced for both slow speed conditions (Slow-Far; *t* =-2.74, *p=*0.007, Slow-Close; *t* = -2.60 *p*=0.01) but not for either fast speed condition (Fast-Far; *t* =-0.39, *p*=0.701, Fast-Close; t = -0.47, *p*=0.641), suggesting that suppression starts earlier when the target is moving slower.

To further examine the interaction between target speed and time of TMS delivery for the FDI muscle, we performed a growth curve analysis (Wiley 2014) averaged across the levels of target speed (Figure 6A; see *Statistical Analysis* for details). Here we found a significant effect of target speed on the intercept term, indicating lower overall MEP amplitude proportions for the Slow condition relative to the Fast condition (Estimate= -4.333, SE= 1.99, *p*=0.02). There was also a significant effect on the quadratic term, indicating shallower curvature in the slow condition relative to the fast condition (Estimate= -1.003, SE= 5.09, *p*=0.008). All other effects of target speed were not significant.

**Figure 6.**
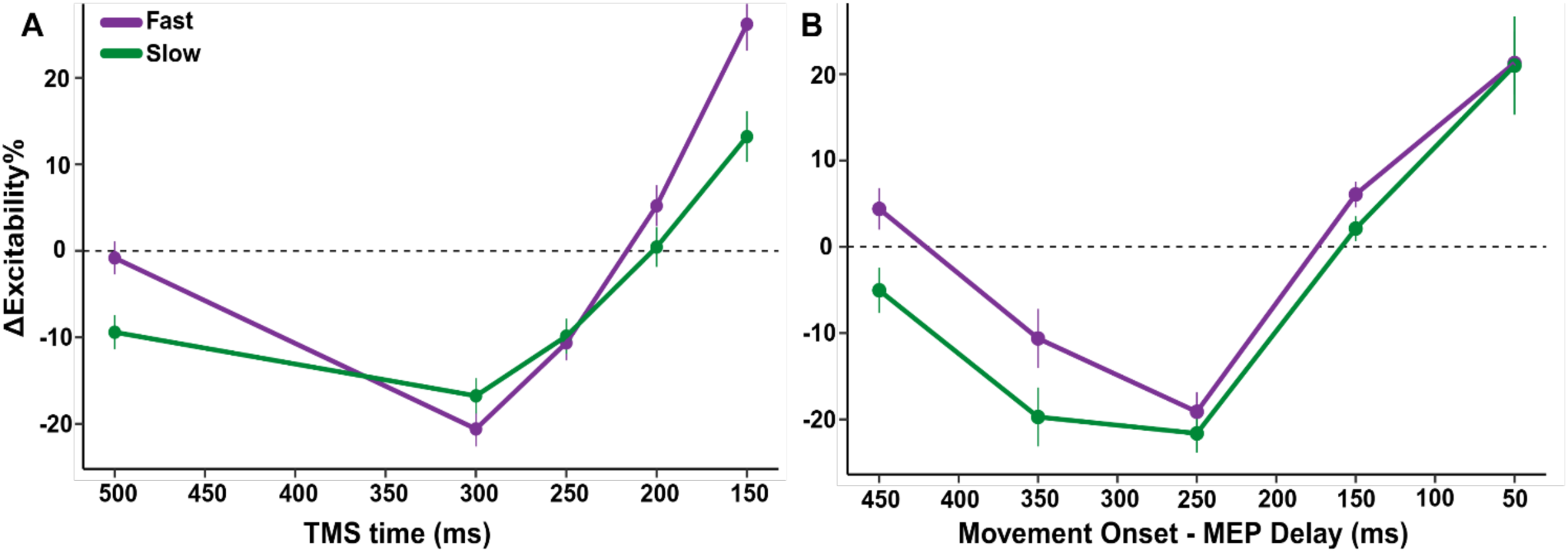
Increased overall excitability for faster-moving targets when adjusting for movement onset. Growth curve analysis for FDI excitability between speed conditions relative to (**A**) TMS time and (**B**) movement onset. Error bars correspond to 95% confidence interval.

Given that behaviorally, we observed earlier movement onsets for faster-moving targets (see Figures 2 and 3), we conducted a second growth curve analysis in which we computed the difference between the time of TMS delivery and movement onset so that MEPs would be aligned to the EMG-based time of movement onset instead of the target’s arrival (Figure 6B). Trials were sorted into 100 ms time bins centered around 450, 350, 250, 150, and 50 ms before movement. There was still a significant effect of target speed on the intercept term, indicating overall higher levels of CSE for fast targets relative to slow targets (Estimate = -5.78, SE= 2.74, *p*=0.04). However, there was no longer a significant effect of speed on the quadratic term, indicating that the curvature of preparatory suppression was similar between slow and fast trials (Estimate= -6.50, SE= 6.4, *p*= 0.32). All other effects of target speed were not significant. These results suggest that, though faster target speeds heighten CSE overall, the specific pattern of CSE modulation is relative to response execution.

In the task-irrelevant muscle (ADM; grey lines in Figure 5), there were significant main effects of target distance (*F*_1,21_=5.20, η^2^=0.005, *p*=0.03) and time of TMS (*F*_2.21, 46.48_=7.33, η^2^=.044, *p*=0.001), but not for target speed (*F*_1,21_=4.22, η^2^=0.006, *p*=0.053). In addition, a significant two-way interaction between target distance and TMS delivery time for CSE in the ADM muscle was observed (*F*_3.31,69.56_=2.69, η^2^=0.010, *p*=0.048), indicating that ADM showed greater early suppression at farther starting distances (see Figure 5). Post-hoc tests showed that ADM MEP amplitudes at 500 ms and 300 ms before the target’s arrival were significantly reduced for the Slow-Close, Slow-Far, and Fast-Far (all *p’*s<0.001) conditions, but not for Fast-Close (500 ms: *t* =0.145, *p*=0.885; 300 ms: *t* = -1.28, *p*=0.201) conditions. These results suggest that the early suppression of task-irrelevant muscles is apparent only with longer (>750 ms) preparatory periods.

## Discussion

In the present study, we examined how target speed and distance influence TMS-evoked motor excitability during the preparation phase of manual interceptions. We hypothesized that if M1 integrates kinematic information to estimate TTC, then CSE modulation would vary depending on both target speed and distance, and this modulation would be associated with interception timing. In partial support of our hypothesis, we found that the time course of this CSE modulation was influenced by target speed, but not by its starting distance. Notably, after adjusting for movement onset, we observed reduced preparatory suppression and a general increase in excitability for faster-moving targets. This pattern may reflect an enhanced urgency signal that contributes to earlier movement initiation, reduced movement time, and decreased spatial accuracy with increasing target speed. Overall, our results suggest that target visual motion properties modulate corticospinal excitability, playing a crucial role in shaping movement preparation during interception.

Our findings further support that CSE modulation follows a distinct timecourse of suppression and facilitation during movement preparation. As expected, the timing of both suppression and facilitation in this task was tightly linked with movement onset, with significant MEP suppression relative to baseline observed approximately 250-300 ms before EMG onset (Duque et al. 2017; Duque and Ivry 2009; Hannah et al. 2018; Hasbroucq et al. 1997; Lebon et al. 2016; Touge et al. 1998). As movement onset approached (∼150-50 ms prior), there was a significant increase in CSE in the task-relevant muscle (FDI), consistent with previous studies showing a buildup of excitability immediately before movement execution (Chen et al. 1998; Coxon et al. 2006; Leocani et al. 2000; MacKinnon and Rothwell 2000). This facilitatory effect coincides with increased activity measured in primate motor neurons during preparation (Chen and Hallett 1999; Port et al. 2001). In contrast, the task-irrelevant muscle (ADM) exhibited increasing suppression throughout the preparation phase, suggesting an inhibitory mechanism that optimizes motor output by selectively suppressing unnecessary muscles (Greenhouse et al. 2015; Hannah et al. 2018; Labruna et al. 2019; Quoilin et al. 2019). Collectively, our results extend previous work, demonstrating a similar pattern of corticospinal suppression and facilitation during the preparation of visually guided interceptive movements.

Unlike delayed RT tasks, where movement initiation follows an unpredictable cue, the present task allows for active preparation while the target is in motion. Our finding of suppression in this task would therefore go against the impulse control theory of preparatory suppression (Duque et al. 2010; Duque and Ivry 2009; Greenhouse et al. 2015; Vassiliadis et al. 2020), as withholding a prepared response is not required in our task. Instead, our results support the idea that inhibitory mechanisms within the M1 enable effective movement preparation (Hannah et al. 2018; Ibáñez et al. 2020). The selective suppression and rapid release of inhibition in task-relevant muscles just before execution may also reflect a surround inhibition or gain modulation mechanism, such that inhibition serves to sculpt corticospinal output underlying precise motor control (Beck and Hallett 2011; Duque et al. 2017; Greenhouse 2022; Lebon et al. 2016). In the case of interception, this modulation of inhibitory processes likely aids in integrating visual motion information and informs movement timing (Fujiyama et al. 2012). Altogether, our findings underscore the importance of considering context-specific and dynamic task environments to advance our understanding of the mechanisms underlying preparatory CSE modulation.

Our results suggest that target speed-dependent CSE modulation underlies earlier movement initiation when trying to intercept faster moving targets. To our knowledge, no previous study has measured CSE in scenarios where accurately timing movement onset depends on using visual information to estimate the target’s time-to-contact (TTC) on a trial-by-trial basis. Our approach allowed us to distinguish between the relative influence of target speed and distance, while accounting for the duration of the preparatory phase. Interestingly, though the overall pattern of CSE modulation was similar for both fast and slow-moving targets, faster targets resulted in less suppression and increased facilitation. Increased CSE facilitation for faster-moving targets may reflect M1 reaching an excitatory threshold sooner, causing an earlier release of motor commands (Chen et al. 1998; Hanes and Schall 1996). Alternatively, this earlier transition from suppression to facilitation could represent an adjustment to the heightened demands on accuracy when intercepting faster targets (Brouwer et al. 2005; Tresilian and Lonergan 2002). Additionally, we observed early preparatory suppression at our earliest tested timepoint (TMS_500_) only for slow targets, whereas suppression for fast targets relative to baseline only appeared at later timepoints (TMS_300_ and TMS_250_). This finding is consistent with previous studies showing that shortened preparatory periods (e.g., 350 ms) can eliminate motor-evoked potential (MEP) suppression, while longer preparation times (e.g., 1400 ms) maintain suppression (McInnes et al. 2021). Thus, it is possible that preparatory suppression is not a universal feature of all movements but rather indicates an adaptive mechanism to optimize response timing. In the present study, when preparing for fast-moving targets, the motor system may prioritize a quicker release of the motor command, enabling timely interception under more urgent conditions (Ball et al. 1999; Fooken et al. 2024; McInnes et al. 2021).

When adjusting for movement onset times, the pattern of CSE modulation was comparable between fast and slow targets, but there was an overall heightened excitability for fast targets. This suggests that CSE modulation is more closely associated with the initiation of movement rather than the underlying processes of movement preparation (Haith et al. 2016). This earlier onset and continued elevated CSE align with the idea of increased response urgency (Carland et al. 2019; Cisek et al. 2009; McInnes et al. 2023; Thura 2020) for intercepting fast-moving targets (Barany et al. 2020; Gómez-Granados et al. 2021), resulting in quicker response initiation (Brouwer et al. 2005). The urgency-related CSE modulation observed in the present study may be partially due to the overall shorter duration of preparation when the target is moving faster (McInnes et al. 2021). Nonetheless, we observed some evidence of reduced early suppression and increased late facilitation between fast and slow targets even when the preparation duration was identical, mirroring behavioral differences in movement onset (compare Fast-Far vs. Slow-Close conditions in Figures 2 and 5). Together, these results suggest that target speed modulates the urgency signal, driving increased excitability for intercepting faster moving targets (Derosiere et al. 2022).

While we used TMS primarily to examine changes in CSE during interception preparation, we also observed that TMS applied close to target arrival led to delayed movement onset (Figure 3A), consistent with previous research indicating that TMS can delay RT when applied near the expected response time (Leocani et al. 2000; Pascual-Leone et al. 1992; Ziemann et al. 1997). This delay is typically attributed to the cortical silent period, which temporarily suppresses M1 neuron activity (Leocani et al. 2000; Ziemann et al. 1997). Notably, participants compensated for this delay by shortening their total movement time (compare Figure 3A and 3B), suggesting that TMS may interfere with motor initiation without affecting online motor control. Interestingly, when TMS was applied 500 ms before target arrival, movement occurred significantly earlier compared to conditions without TMS for the three conditions with longer preparation durations. The earlier movement initiation could be attributed to intersensory facilitation, in which the sensory input from the TMS pulse enhances the processing of the imperative stimulus (in this case integrating visual information about the moving target), thereby speeding up movement onset (Ibáñez et al. 2020). This effect was not observed in the Fast-Close condition, likely due to the already constrained movement onset imposed by the short motion duration (750 ms). Further research using dynamic movement execution tasks is needed to clarify the specific mechanisms by which TMS influences motor performance outcomes (Gomez et al. 2022).

## Conclusion

The present study shows that dynamic changes in M1 excitability during preparation play a critical role in estimating TTC for accurate interceptive performance, and that alterations in target speed may modulate both the timing and urgency of motor planning and execution. We found that when preparing to intercept a moving target, CSE undergoes a similar transition profile from early suppression to late facilitation invariant of the speed or distance of the target. However, faster-moving targets led to earlier modulation and heightened excitability, reflecting an increased urgency to respond. In slower target conditions, corticospinal excitability suppression was evident at 500 ms before the ideal interception time, suggesting increased inhibitory mechanisms to optimize timing. The different CSE patterns for fast and slow targets may underline differences in behavioral performance, such that faster-moving targets elicit a faster release of the motor command for earlier movement onset. Furthermore, applying TMS during preparation delayed movement onsets when delivered close to the expected movement onset and sped up onsets when delivered early. Together, these findings underscore the importance of CSE modulation supporting interceptive actions and highlight how variations in target speed impact motor planning and execution.

## Disclosures

Authors report no conflict of interest.

## Acknowledgments

We thank Asher Khan and Emma Whitman for assistance with data collection. This work was supported by the University of Georgia Mary Frances Early College of Education and University of Georgia Office of Research. All figures were created with BioRender (https://BioRender.com/j37f899).

## Notes

https://BioRender.com/j37f899

### Competing Interest Statement

The authors have declared no competing interest.

## References

Badry R, Mima T, Aso T, Nakatsuka M, Abe M, Fathi D, Foly N, Nagiub H, Nagamine T, Fukuyama H. Suppression of human cortico-motoneuronal excitability during the Stop-signal task. Clinical Neurophysiology 120: 1717–1723, 2009.

Ball T, Schreiber A, Feige B, Wagner M, Lücking CH, Kristeva-Feige R. The Role of Higher-Order Motor Areas in Voluntary Movement as Revealed by High-Resolution EEG and fMRI. NeuroImage 10: 682–694, 1999.

Barany DA, Gómez-Granados A, Schrayer M, Cutts SA, Singh T. Perceptual decisions about object shape bias visuomotor coordination during rapid interception movements. Journal of neurophysiology 123: 2235–2248, 2020.

Battaglia-Mayer A, Caminiti R. Parieto-frontal networks for eye–hand coordination and movements. In: Handbook of Clinical Neurology. Elsevier, p. 499–524.

Baurès R, Fourteau M, Thébault S, Gazard C, Pasquio L, Meneghini G, Perrin J, Rosito M, Durand J, Roux F. Time-to-contact perception in the brain. J of Neuroscience Research 99: 455–466, 2021.

Beck S, Hallett M. Surround inhibition in the motor system. Exp Brain Res 210: 165–172, 2011.

Bestmann S, Duque J. Transcranial Magnetic Stimulation: Decomposing the Processes Underlying Action Preparation. Neuroscientist 22: 392–405, 2016.

Bosco G, Carrozzo M, Lacquaniti F. Contributions of the Human Temporoparietal Junction and MT/V5+ to the Timing of Interception Revealed by Transcranial Magnetic Stimulation. J Neurosci 28: 12071–12084, 2008.

Brenner E, Smeets JBJ. Continuously updating one’s predictions underlies successful interception. Journal of Neurophysiology 120: 3257–3274, 2018.

Brouwer A-M, Brenner E, Smeets JBJ. Hitting moving objects. Experimental Brain Research 133: 242–248, 2000.

Brouwer A-M, Middelburg T, Smeets JBJ, Brenner E. Hitting moving targets. Experimental Brain Research 152: 368–375, 2003.

Brouwer A-M, Smeets JBJ, Brenner E. Hitting moving targets: effects of target speed and dimensions on movement time. Exp Brain Res 165: 28–36, 2005.

Carland MA, Thura D, Cisek P. The Urge to Decide and Act: Implications for Brain Function and Dysfunction. Neuroscientist 25: 491–511, 2019.

Chang C-J, Jazayeri M. Integration of speed and time for estimating time to contact. Proc Natl Acad Sci USA 115, 2018.

Chen R, Hallett M. The Time Course of Changes in Motor Cortex Excitability Associated with Voluntary Movement. Can j neurol sci 26: 163–169, 1999.

Chen R, Yaseen Z, Cohen LG, Hallett M. Time course of corticospinal excitability in reaction time and self-paced movements. Annals of Neurology 44: 317–325, 1998.

Cisek P, Puskas GA, El-Murr S. Decisions in Changing Conditions: The Urgency-Gating Model. J Neurosci 29: 11560–11571, 2009.

Coxon JP, Stinear CM, Byblow WD. Intracortical Inhibition During Volitional Inhibition of Prepared Action. Journal of Neurophysiology 95: 3371–3383, 2006.

Crowe EM, Smeets JBJ, Brenner E. Online updating of obstacle positions when intercepting a virtual target. Exp Brain Res 241: 1811–1820, 2023.

Daneshi A, Azarnoush H, Towhidkhah F, Bernardin D, Faubert J. Brain activity during time to contact estimation: an EEG study. Cogn Neurodyn 14: 155–168, 2020.

Darling WG, Wolf SL, Butler AJ. Variability of motor potentials evoked by transcranial magnetic stimulation depends on muscle activation. Exp Brain Res 174: 376–385, 2006.

De Azevedo Neto RM, Bartels A. Disrupting short-term memory in premotor cortex affects serial dependence in visuomotor integration. 2021.

De La Malla C, López-Moliner J. Hitting moving targets with a continuously changing temporal window. Exp Brain Res 233: 2507–2515, 2015.

Derosiere G, Thura D, Cisek P, Duque J. Hasty sensorimotor decisions rely on an overlap of broad and selective changes in motor activity. PLoS Biol 20: e3001598, 2022.

Derosiere G, Vassiliadis P, Duque J. Advanced TMS approaches to probe corticospinal excitability during action preparation. NeuroImage 213: 116746, 2020.

Dubrowski A, Lam J, Carnahan H. Target velocity effects on manual interception kinematics. Acta Psychologica 104: 103–118, 2000.

Duque J, Greenhouse I, Labruna L, Ivry RB. Physiological Markers of Motor Inhibition during Human Behavior. Trends in Neurosciences 40: 219–236, 2017.

Duque J, Ivry RB. Role of Corticospinal Suppression during Motor Preparation. Cerebral Cortex 19: 2013–2024, 2009.

Duque J, Lew D, Mazzocchio R, Olivier E, Ivry RB. Evidence for Two Concurrent Inhibitory Mechanisms during Response Preparation. J Neurosci 30: 3793–3802, 2010.

Field DT, Wann JP. Perceiving Time to Collision Activates the Sensorimotor Cortex. Current Biology 15: 453–458, 2005.

Fooken J, Balalaie P, Park K, Flanagan JR, Scott SH. Rapid eye and hand responses in an interception task are differentially modulated by context-dependent predictability. Journal of Vision 24: 10, 2024.

Fooken J, Baltaretu BR, Barany DA, Diaz G, Semrau JA, Singh T, Crawford JD. Perceptual-Cognitive Integration for Goal-Directed Action in Naturalistic Environments. J Neurosci 43: 7511–7522, 2023.

Fujiyama H, Hinder MR, Schmidt MW, Tandonnet C, Garry MI, Summers JJ. Age-related Differences in Corticomotor Excitability and Inhibitory Processes during a Visuomotor RT Task. Journal of Cognitive Neuroscience 24: 1253–1263, 2012.

Gallivan JP, Culham JC. Neural coding within human brain areas involved in actions. Current Opinion in Neurobiology 33: 141–149, 2015.

Gomez IN, Orsinger SR, Kim HE, Greenhouse I. Assessing Corticospinal Excitability During Goal-Directed Reaching Behavior. JoVE 64238, 2022.

Gómez-Granados A, Barany DA, Schrayer M, Kurtzer I, Bonnet C, Singh T. Age-related deficits in rapid visuomotor decision-making. 2021.

Greenhouse I. Inhibition for gain modulation in the motor system. Exp Brain Res 240: 1295–1302, 2022.

Greenhouse I, Sias A, Labruna L, Ivry RB. Nonspecific Inhibition of the Motor System during Response Preparation. Journal of Neuroscience 35: 10675–10684, 2015.

Haith AM, Bestmann S. Preparation of Movement. In: The Cognitive Neurosciences, edited by Poeppel D, Mangun GR, Gazzaniga MS. The MIT Press, p. 541–548.

Haith AM, Pakpoor J, Krakauer JW. Independence of Movement Preparation and Movement Initiation. J Neurosci 36: 3007–3015, 2016.

Hanes DP, Schall JD. Neural Control of Voluntary Movement Initiation. Science 274: 427–430, 1996.

Hannah R, Cavanagh SE, Tremblay S, Simeoni S, Rothwell JC. Selective Suppression of Local Interneuron Circuits in Human Motor Cortex Contributes to Movement Preparation. J Neurosci 38: 1264–1276, 2018.

Hasbroucq T, Kaneko H, Akamatsu M, Possamaї C-A. Preparatory inhibition of cortico-spinal excitability: a transcranial magnetic stimulation study in man. Cognitive Brain Research 5: 185–192, 1997.

Holm S. A Simple Sequentially Rejective Multiple Test Procedure. Scandinavian Journal of Statistics 6: 65–70, 1979.

Ibáñez J, Hannah R, Rocchi L, Rothwell JC. Premovement Suppression of Corticospinal Excitability may be a Necessary Part of Movement Preparation. Cerebral Cortex 30: 2910–2923, 2020.

Jackson N, Greenhouse I. VETA: An Open-Source Matlab-Based Toolbox for the Collection and Analysis of Electromyography Combined With Transcranial Magnetic Stimulation. Front Neurosci 13: 975, 2019.

Labruna L, Lebon F, Duque J, Klein P-A, Cazares C, Ivry RB. Generic Inhibition of the Selected Movement and Constrained Inhibition of Nonselected Movements during Response Preparation. Journal of Cognitive Neuroscience 26: 269–278, 2014.

Labruna L, Tischler C, Cazares C, Greenhouse I, Duque J, Lebon F, Ivry RB. Planning face, hand, and leg movements: anatomical constraints on preparatory inhibition. Journal of Neurophysiology 121: 1609–1620, 2019.

Lebon F, Greenhouse I, Labruna L, Vanderschelden B, Papaxanthis C, Ivry RB. Influence of Delay Period Duration on Inhibitory Processes for Response Preparation. Cereb Cortex 26: 2461–2470, 2016.

Lee DN. A Theory of Visual Control of Braking Based on Information about Time-to-Collision. Perception 5: 437–459, 1976.

Leocani L, Cohen LG, Wassermann EM, Ikoma K, Hallett M. Human corticospinal excitability evaluated with transcranial magnetic stimulation during different reaction time paradigms. Brain 123: 1161–1173, 2000.

Li Y, Mo L, Chen Q. Differential contribution of velocity and distance to time estimation during self-initiated time-to-collision judgment. Neuropsychologia 73: 35–47, 2015.

MacKinnon CD, Rothwell JC. Time-varying changes in corticospinal excitability accompanying the triphasic EMG pattern in humans. The Journal of Physiology 528: 633–645, 2000.

Marinovic W, Plooy AM, Tresilian JR. The Utilisation of Visual Information in the Control of Rapid Interceptive Actions. Experimental Psychology 56: 265–273, 2009.

Marinovic W, Reid CS, Plooy AM, Riek S, Tresilian JR. Delayed inhibition of an anticipatory action during motion extrapolation. Behav Brain Funct 6: 22, 2010.

McInnes AN, Lipp OV, Tresilian JR, Vallence A, Marinovic W. Premovement inhibition can protect motor actions from interference by response-irrelevant sensory stimulation. The Journal of Physiology 599: 4389–4406, 2021.

McInnes AN, Smithers B, Lipp OV, Tresilian JR, Vallence A-M, Rothwell JC, Marinovic W. From Inhibition to Excitation and Why: The Role of Temporal Urgency in Modulating Corticospinal Activity. 2023.

Merchant H, Zarco W, Prado L, Pérez O. Behavioral and Neurophysiological Aspects of Target Interception. In: Progress in Motor Control, edited by Sternad D. Springer US, p. 201–220.

Nelson JS, Baud-Bovy G, Smeets JBJ, Brenner E. Accuracy of Intercepting Moving Tactile Targets. Perception 48: 685–701, 2019.

Nickerson RS. Intersensory facilitation of reaction time: Energy summation or preparation enhancement? Psychological Review 80: 489–509, 1973.

O’Reilly JX, Mesulam MM, Nobre AC. The Cerebellum Predicts the Timing of Perceptual Events. J Neurosci 28: 2252–2260, 2008.

Pascual-Leone A, Valls-Solé J, Wassermann EM, Brasil-Neto J, Cohen LG, Hallett M. EFFECTS OF FOCAL TRANSCRANIAL MAGNETIC STIMULATION ON SIMPLE REACTION TIME TO ACOUSTIC, VISUAL AND SOMATOSENSORY STIMULI. Brain 115: 1045–1059, 1992.

Peirce J, Gray JR, Simpson S, MacAskill M, Höchenberger R, Sogo H, Kastman E, Lindeløv JK. PsychoPy2: Experiments in behavior made easy. Behav Res 51: 195–203, 2019.

Polti I, Nau M, Kaplan R, Van Wassenhove V, Doeller CF. Rapid encoding of task regularities in the human hippocampus guides sensorimotor timing. eLife 11: e79027, 2022.

Port NL, Kruse W, Lee D, Georgopoulos AP. Motor Cortical Activity during Interception of Moving Targets. Journal of Cognitive Neuroscience 13: 306–318, 2001.

Quoilin C, Fievez F, Duque J. Preparatory inhibition: Impact of choice in reaction time tasks. Neuropsychologia 129: 212–222, 2019.

Rossini PM, Barker AT, Berardelli A, Caramia MD, Caruso G, Cracco RQ, Dimitrijević MR, Hallett M, Katayama Y, Lücking CH, Maertens De Noordhout AL, Marsden CD, Murray NMF, Rothwell JC, Swash M, Tomberg C. Non-invasive electrical and magnetic stimulation of the brain, spinal cord and roots: basic principles and procedures for routine clinical application. Report of an IFCN committee. Electroencephalography and Clinical Neurophysiology 91: 79–92, 1994.

Satterthwaite FE. An Approximate Distribution of Estimates of Variance Components. Biometrics Bulletin 2: 110, 1946.

Senot P, Baillet S, Renault B, Berthoz A. Cortical Dynamics of Anticipatory Mechanisms in Interception: A Neuromagnetic Study. Journal of Cognitive Neuroscience 20: 1827–1838, 2008.

Senot P, Prévost P, McIntyre J. Estimating time to contact and impact velocity when catching an accelerating object with the hand. Journal of Experimental Psychology: Human Perception and Performance 29: 219–237, 2003.

Sohn YoungH, Hallett M. Surround inhibition in human motor system. Exp Brain Res 158, 2004.

Thura D. Decision urgency invigorates movement in humans. Behavioural Brain Research 382: 112477, 2020.

Touge T, Taylor JL, Rothwell JC. Reduced excitability of the cortico-spinal system during the warning period of a reaction time task. Electroencephalography and Clinical Neurophysiology/Electromyography and Motor Control 109: 489–495, 1998.

Tran DMD, McNair NA, Harris JA, Livesey EJ. Expected TMS excites the motor system less effectively than unexpected stimulation. NeuroImage 226: 117541, 2021.

Tresilian J, Lonergan A. Intercepting a moving target: effects of temporal precision constraints and movement amplitude. Experimental Brain Research 142: 193–207, 2002.

Vassiliadis P, Derosiere G, Grandjean J, Duque J. Motor training strengthens corticospinal suppression during movement preparation. Journal of Neurophysiology 124: 1656–1666, 2020.

Wiley J. Growth Curve Analysis and Visualization Using *R*. J Stat Soft 58, 2014.

Zhao H, Straub D, Rothkopf CA. The visual control of interceptive steering: How do people steer a car to intercept a moving target? Journal of Vision 19: 11, 2019.

Ziemann U, Tergau F, Netz J, Hömberg V. Delay in simple reaction time after focal transcranial magnetic stimulation of the human brain occurs at the final motor output stage. Brain Research 744: 32–40, 1997.

